# Establishing Primary and Stable Cell Lines from Frozen Wing Biopsies for Cellular, Physiological, and Genetic Studies in Bats

**DOI:** 10.1101/2024.03.22.586286

**Authors:** Fengyan Deng, Pedro Morales-Sosa, Andrea Bernal-Rivera, Yan Wang, Dai Tsuchiya, Jose Emmanuel Javier, Nicolas Rohner, Chongbei Zhao, Jasmin Camacho

## Abstract

Bats stand out among mammalian species for their exceptional traits, including the capacity to navigate through flight and echolocation, conserve energy through torpor/hibernation, harbor a multitude of viruses, exhibit resistance to disease, survive harsh environmental conditions, and demonstrate exceptional longevity compared to other mammals of similar size. *In vivo* studies of bats can be challenging for several reasons such as ability to locate and capture them in their natural environments, limited accessibility, low sample size, environmental variation, long lifespans, slow reproductive rates, zoonotic disease risks, species protection, and ethical concerns. Thus, establishing alternative laboratory models is crucial for investigating the diverse physiological adaptations observed in bats. Obtaining quality cells from tissues is a critical first step for successful primary cell derivation. However, it is often impractical to collect fresh tissue and process the samples immediately for cell culture due to the resources required for isolating and expanding cells. As a result, frozen tissue is typically the starting resource for bat primary cell derivation. Yet, cells in frozen tissue are usually damaged and represent low integrity and viability. As a result, isolating primary cells from frozen tissues poses a significant challenge. Herein, we present a successfully developed protocol for isolating primary dermal fibroblasts from frozen bat wing biopsies. This protocol marks a significant milestone, as this the first protocol specially focused on fibroblasts isolation from bat frozen tissue. We also describe methods for primary cell characterization, genetic manipulation of primary cells through lentivirus transduction, and the development of stable cell lines.

**Basic Protocol 1:** Bat wing biopsy collection and preservation

**Support Protocol 1:** Blood collection from bat-venipuncture

**Basic Protocol 2:** Isolation of primary fibroblasts from adult bat frozen wing biopsy

**Support Protocol 2:** Maintenance of primary fibroblasts

**Support Protocol 3:** Cell banking and thawing of primary fibroblasts

**Support Protocol 4:** Growth curve and doubling time

**Support Protocol 5:** Lentiviral transduction of bat primary fibroblasts

**Basic Protocol 3:** Bat stable fibroblasts cell lines development

**Support Protocol 6:** Bat fibroblasts validation by immunofluorescence staining

**Support Protocol 7:** Chromosome counting

## INTRODUCTION

Bats, which make up one fifth of all mammals, have become an important experimental model due to their unique biological features [1–8]. They are the only active flying mammals and can use laryngeal echolocation to orient in darkness [8]. Bats are distributed across every continent except Antarctica [9] with a diverse habitat from tropical rainforests to boreal forests, arid regions and oceanic islands [10]. They perform essential ecological functions that contribute to the health of our environment, including pest control, seed dispersal, and pollination [11]. Bats have also been identified as natural reservoirs of many infectious disease agents including Ebola virus [12, 13], Marburg virus [13], SARS-like Coronaviruses [14, 15], Middle East respiratory syndrome coronavirus (MERS-CoV) [16] and so on. Despite the importance of bats in ecosystems and epidemiology, the adaptations of their physiology and how their cells handle different viruses are poorly understood. The establishment of *in vitro* bat cell models will facilitate studies of their physiological adaptations and potential molecular implications to their heightened antiviral defenses [17], tolerance to viral infections [18, 19], ability to survive in harsh conditions [20–23], and longevity [24, 25].

Acquiring high-quality cells from tissues is an essential initial stage for the successful derivation of fibroblasts and other primary cell types. During field work to capture bats, the feasibility of collecting fresh tissue and promptly processing the samples for cell culture is hindered by the laboratory resources required for isolating and expanding cells, including specialized media, drug treatments, 4°C cold storage, centrifuge, microscope, cell incubators, and access to -80°C storage [15]. As a result, frozen tissue is commonly the initial resource for deriving primary bat cells. Obtaining high-quality cells from frozen tissues is a pivotal first step for primary cell derivation. However, frozen tissue often contains damaged cells, leading to reduced integrity and viability, making the derivation of primary cells particularly challenging. To address this, we have developed a rapid, field-ready method to cryopreserve wing biopsies for optimal recovery of healthy cells from frozen skin samples (Fig. 1).

**Figure 1.**
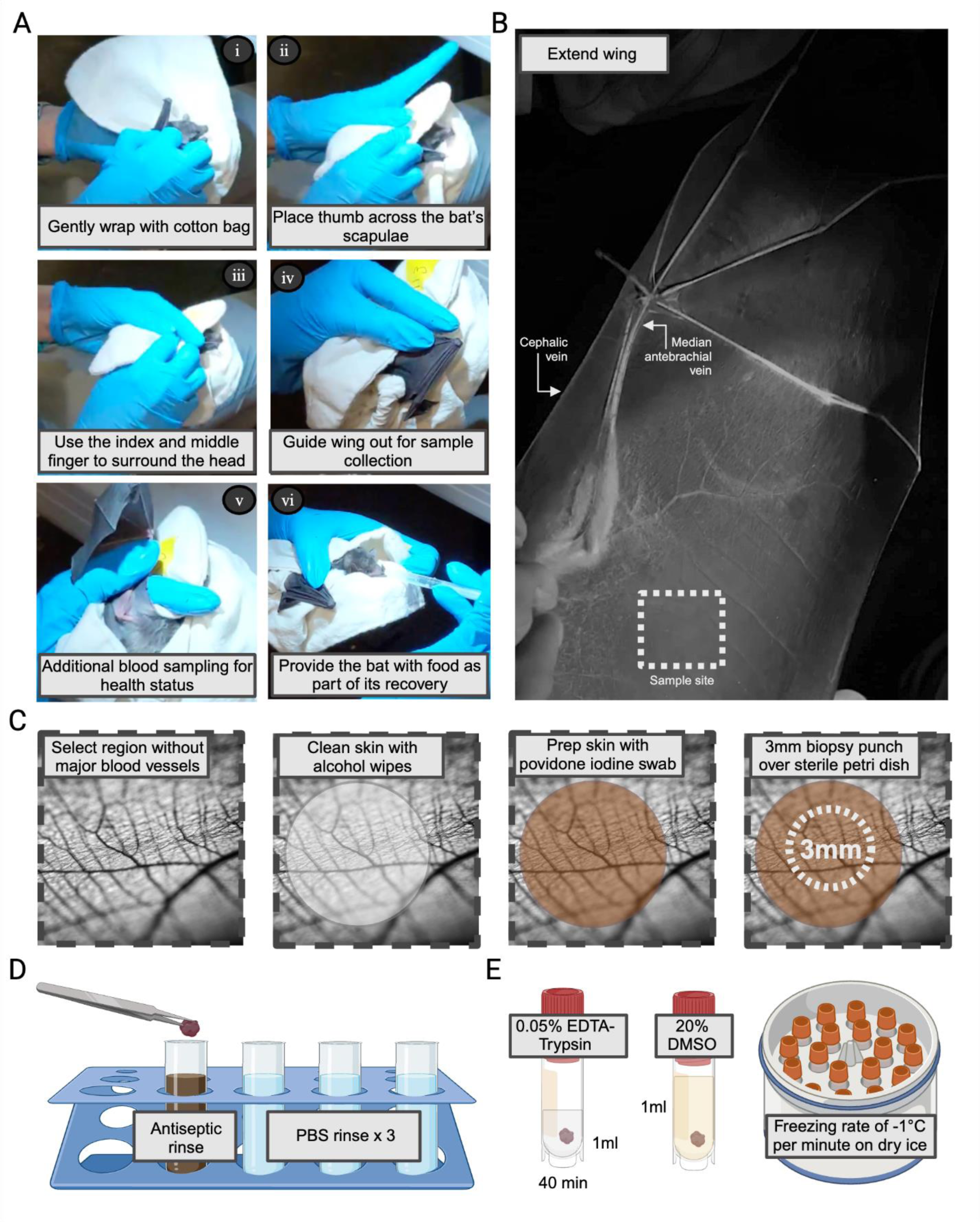
Workflow of bat wing biopsy collection and cryopreservation. This works best with two people but can be done alone. A. The bat is gently wrapped in a cotton bat bag, with head covered, to reduce stress prior to sampling. B. The wing is first outstretched and inspected for a sampling site at the plagiopatagium, close to the body, and free from major blood vessels, muscle fibers, and hair. C. The skin of the sample site is cleaned with 70% ethanol and antiseptic. The dotted circle shows an example of one 3 mm wing biopsy punch taken in an area with little vascularization. D. The biopsy punch was washed in antiseptic and PBS. E. The small bat wing biopsy was digested in EDTA-Trypsin before adding freezing media, followed by cryopreservation.

Fibroblasts are the most common cells in connective tissue and are distributed across every organ system in the body [26]. They are easy to grow and are commonly used to produce induced pluripotent stem cells (iPSC) to generate other cell types [27]. Besides, fibroblasts are the principal source of the extensive extracellular matrix (ECM), and play essential roles in wound healing, inflammation response, tissue regeneration, angiogenesis, cancer progression, and viral infection progression [10]. Thus, fibroblasts are often used to study disease related inflammation [28, 29] and innate immune response across species [11–13]. The isolation of bat fibroblasts and the development of stable fibroblast cell lines would contribute to the understanding of bat biology and its implications for various fields, including ecology, virology, and comparative physiology [11, 12, 14].

The double-layered epidermis of the bat wing (Fig. 2A) provides strength to withstand the stresses of flight and serves as a robust barrier against potential pathogens encountered in caves or during flight. If damage occurs in the wing, it can rapidly heal the wound due to immune cells (Fig. 2B) migrating to the injury site [30, 31], dermal fibroblasts proliferating and producing ECM, followed by remodeling of the newly formed tissue [32]. However, due to the special structure of the bat wing skin tissue (Fig. 1A), normal skin fibroblasts derivation method is not efficient enough for fibroblasts derivation from bat wing tissue. In this article, we have developed a protocol for isolating fibroblasts from frozen bat wing biopsies obtained from adult bats in the wild (Fig. 2C). The isolated fibroblasts can be cultured, expanded, frozen, thawed, assayed, genetically edited, and used for biological experiments. In our hands, fibroblasts (Fig. 3) isolated from cryopreserved wing tissue can be cultured for more than 20 passages and for at least four months, while maintaining similar morphology and growth rate (Fig. 5). We have successfully applied this procedure to wing tissue isolated from two distinct lineages of bats from the families *Mormoopidae* (13 known species) and *Phyllostomidae* (216 known species). Tissues were obtained from two species, *Pteronotus mesoamericanus* Smith, 1972 (*P. Meso*) (16-19.5g) and *Micronycteris schmidtorum* Sanborn 1935 (*M. Sch*) (7-10g) [33]. With cell lines established, we have developed an improved bat fibroblast growth media (Fig. 5, 6), optimized genetic manipulation using lentiviral transduction (Fig. 7), and standardized protocols to characterize bat fibroblasts with molecular markers (Fig. 8) and karyotyping (Fig. 9). Overall, our study enables the easy use of cell isolation and cell expansion from cryopreserved tissue for future research.

**Figure 2.**
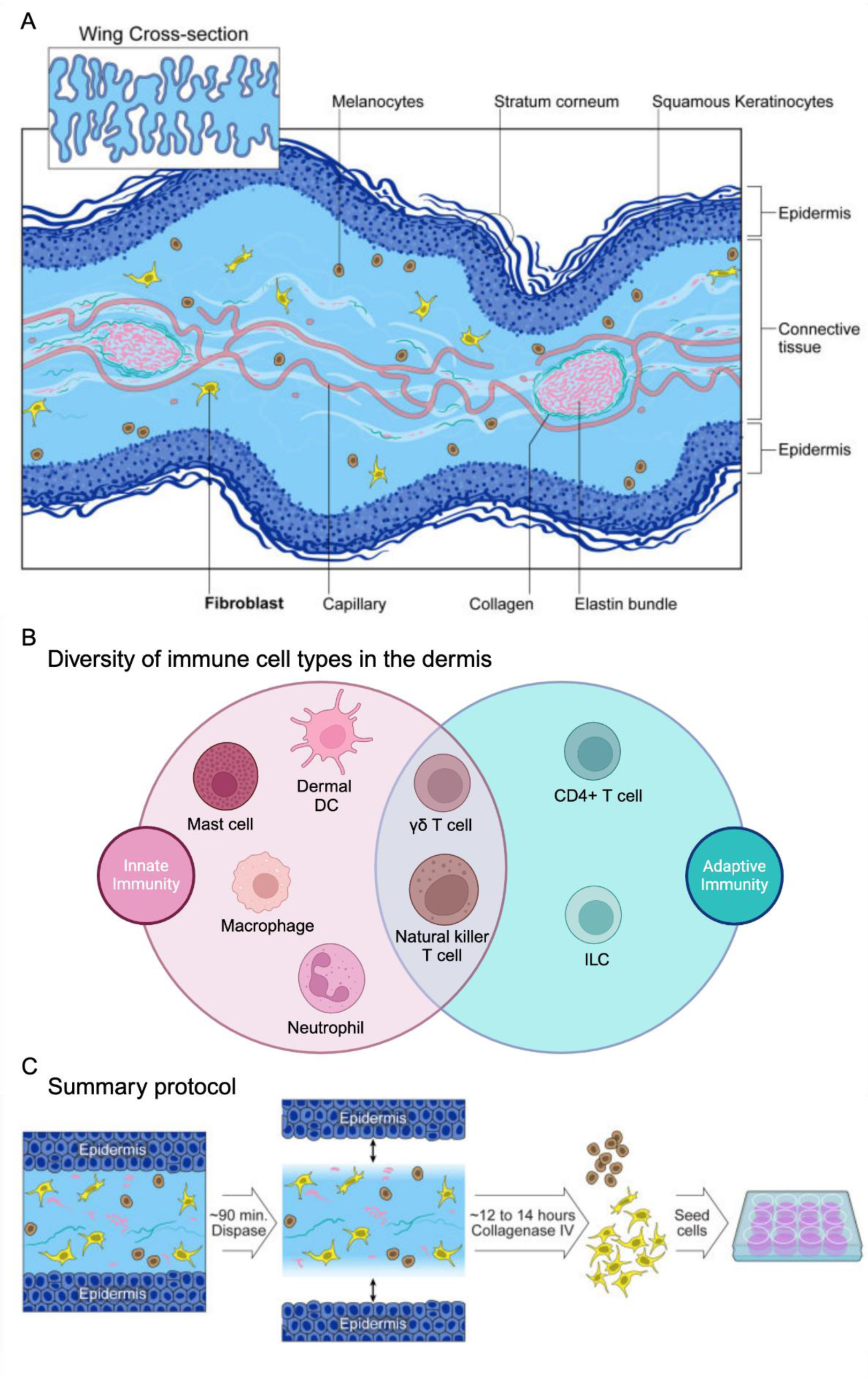
Bat wing layers and workflow of primary fibroblasts isolation. A. Bat wing structure. The bat wing comprised two epidermis and connective tissue layer. The epidermis, which contains stratum corneum and non-keratinized, made up most proportion of the bat wing thickness. The connective tissue layer is composed mostly of elastic and collagen fibres, with capillary perfused in the middle. B. Additional cell types within the dermis. C. Overview of bat wing fibroblasts derivation. Frozen small pieces of bat wing biopsy were dissociated with dispase to expose the connective tissue, which was further digested with collagenase and centrifuged. Cells and small tissue pieces were resuspended with bat primary fibroblasts culture media and seeded on gelatin coated plates.

**Figure 3.**
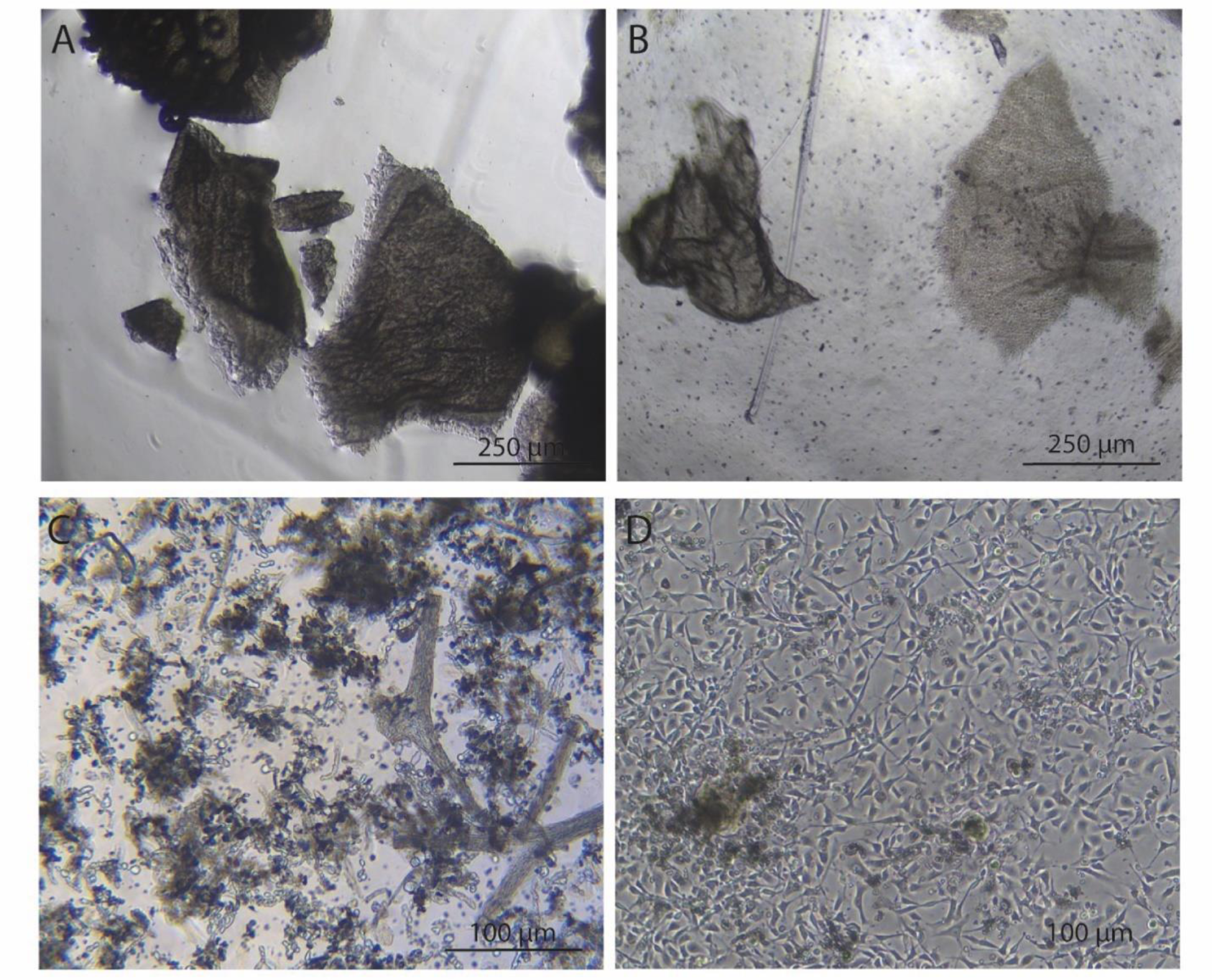
Phase-contrast images showing the morphology of bat wing tissue and cells at different stages of primary fibroblasts isolation. A. bat wing species before dissociation. B. bat wing tissues after dispase dissociation. C. After collagenase dissociation. D. Cells at 5-day post derivation.

**Figure 4.**
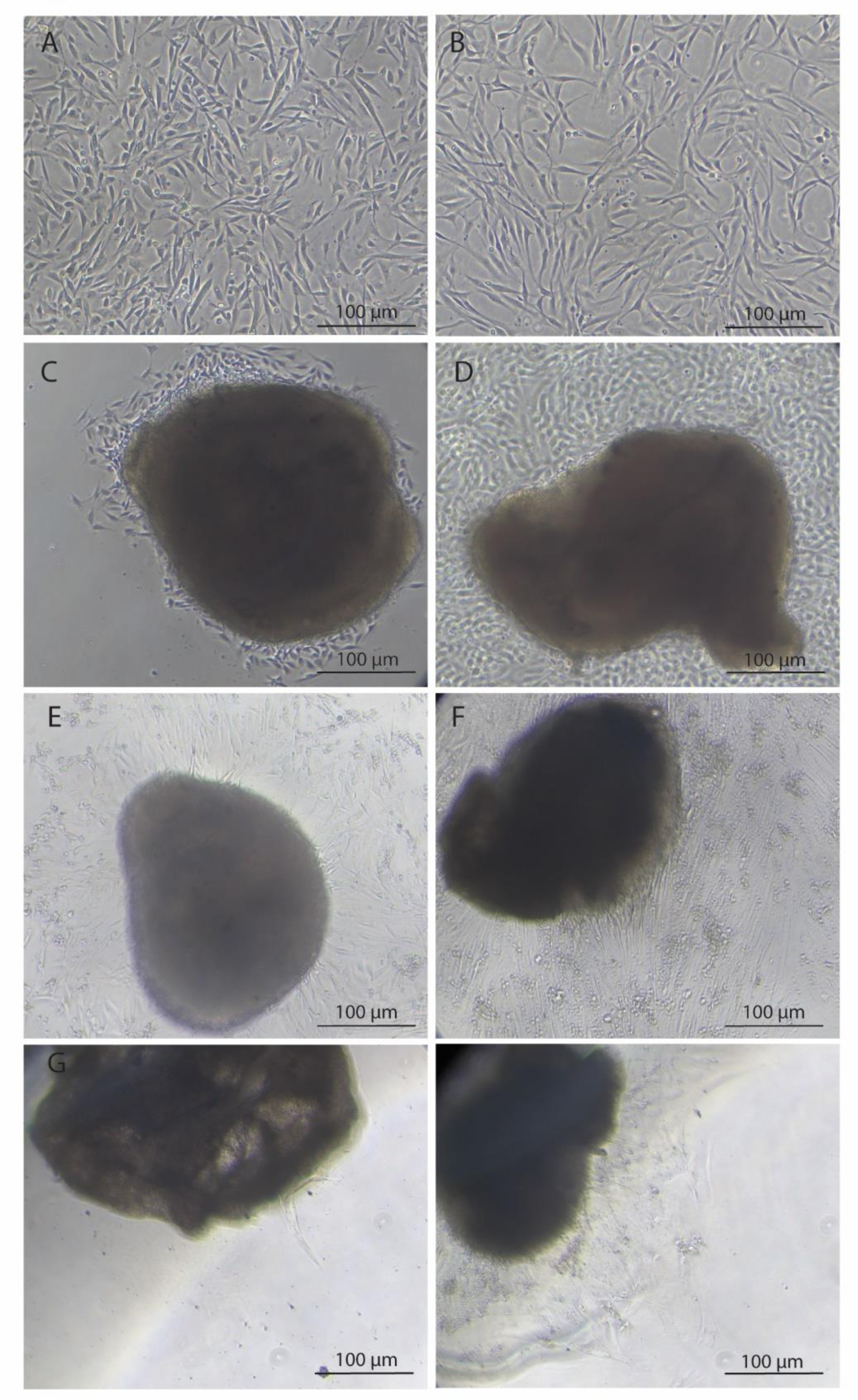
Phase-contrast images showing the morphology of *M. Sch* fibroblasts and bat wing explant from different timepoints. A. *M. Sch* fibroblast cells P1. B. *M. Sch* fibroblast cells P15. C. *M. Sch* wing tissue explant at 17-day post derivation. D. *M. Sch* wing tissue explant at 25-day post derivation. E. *M. Sch* wing tissue explant at 45-day post derivation. F. *M. Sch* wing tissue explant at 98-day post derivation. G. *M. Sch* wing tissue explant at 98-day, 0 day after transferring into new well. H. *M. Sch* wing tissue explant at 103-day post derivation, 5 days after transferring into new well.

**Figure 5.**
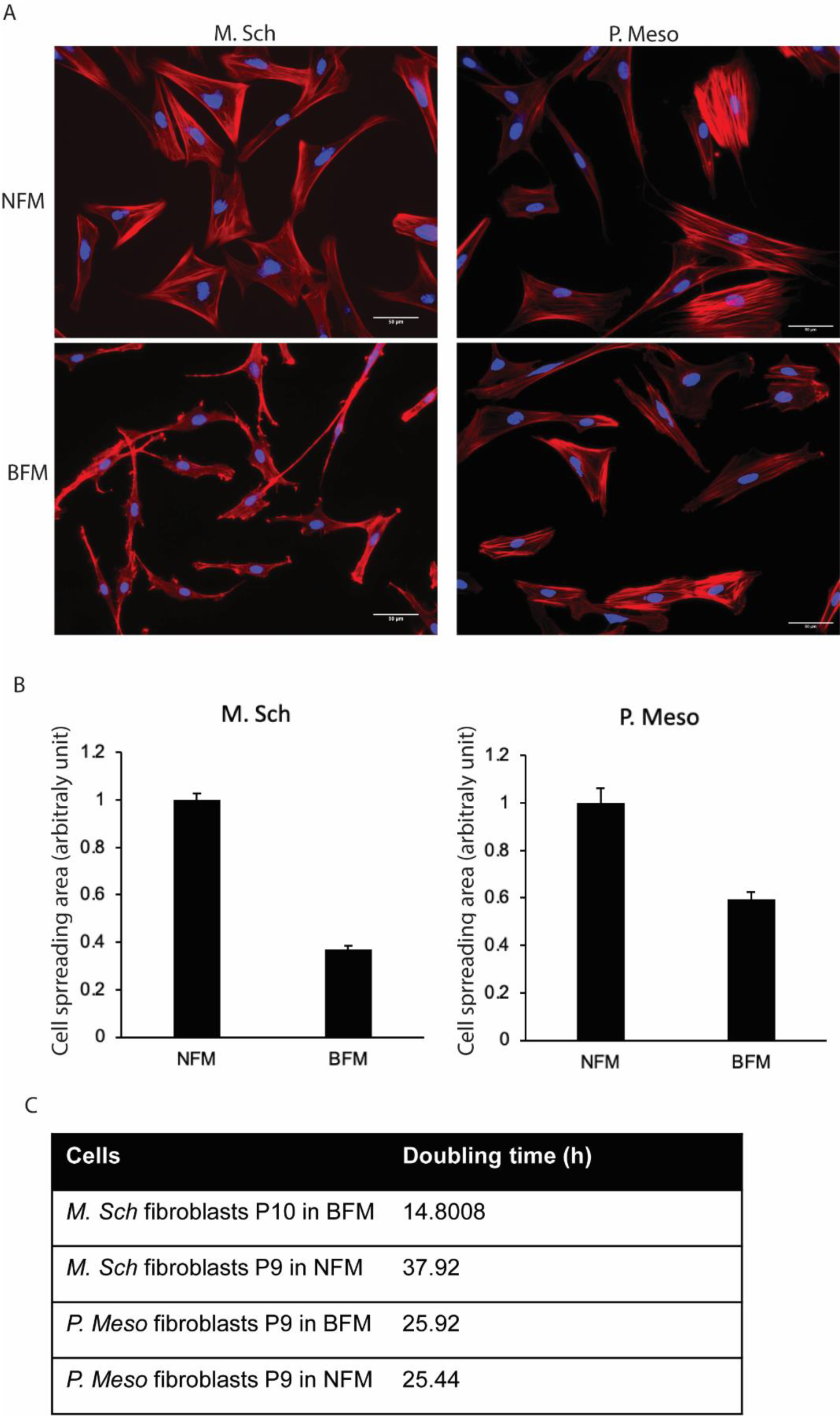
Morphology, size and cell proliferation of fibroblasts during parallel cultivation in normal fibroblasts media (NFM) and bat primary fibroblasts media (BFM). A. Fluorescence microscope images obtained upon F-actin staining of fibroblasts. scale bar = 50 μm; red fluorescence: F-actin; blue fluorescence: DAPI. B. fibroblasts size determined using Image J software. 60 cells from each group were measured for quantification. C. Doubling time of fibroblasts cultured in NFM and BFM. Doubling time of the cells were calculated using online Doubling Time Computing software: https://www.doubling-time.com/compute.php.

**Figure 6.**
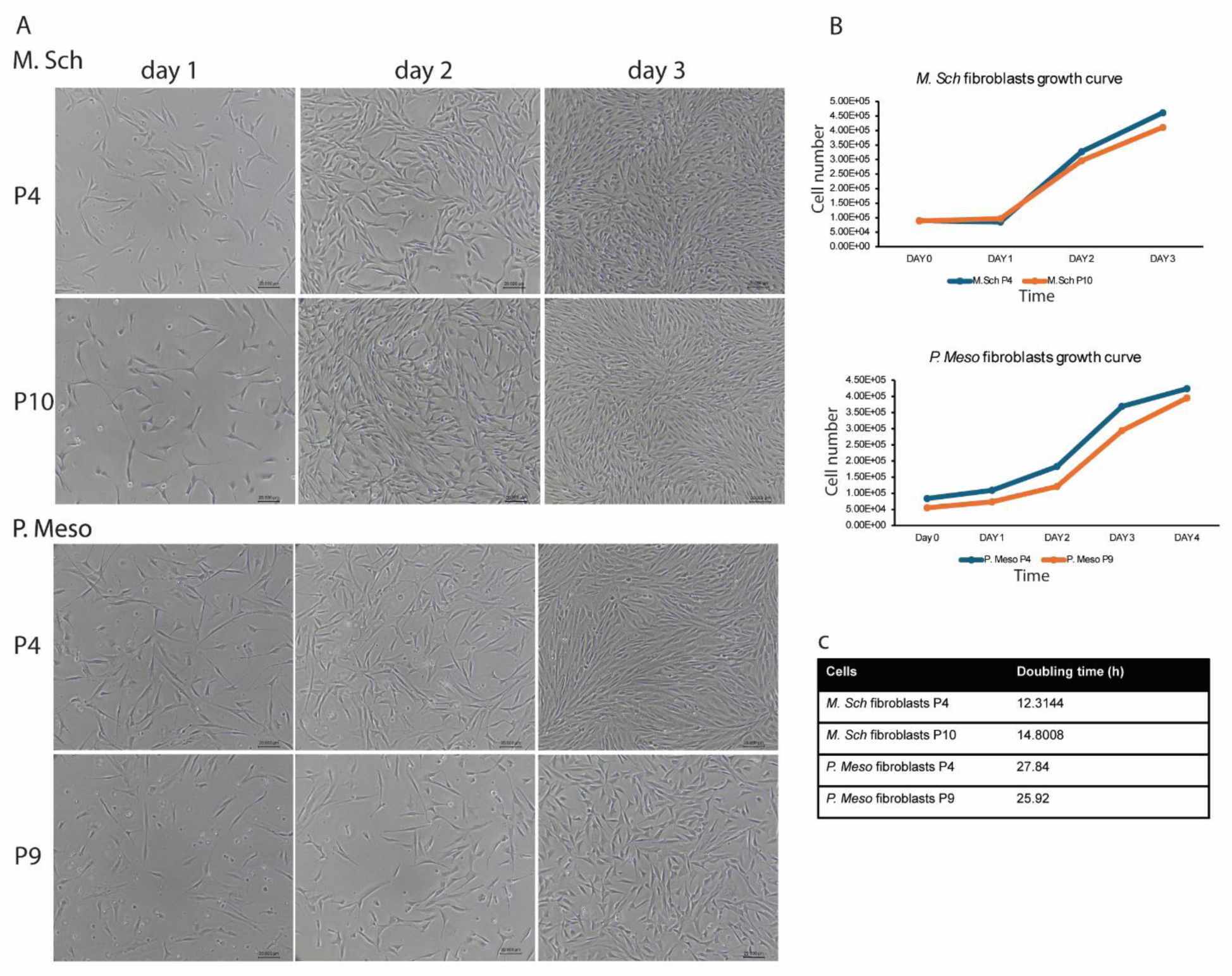
Bat fibroblasts proliferation. A. Phase-contrast images showing the morphology of *M. Sch* and *P. Meso* fibroblasts at different timepoints. B. Growth curve of *M. Sch* and *P. Meso* fibroblasts at different passages. C. doubling time of *M. Sch* and *P. Meso* fibroblasts at different passages. Doubling Time Computing software: https://www.doubling-time.com/compute.php.

**Figure 7.**
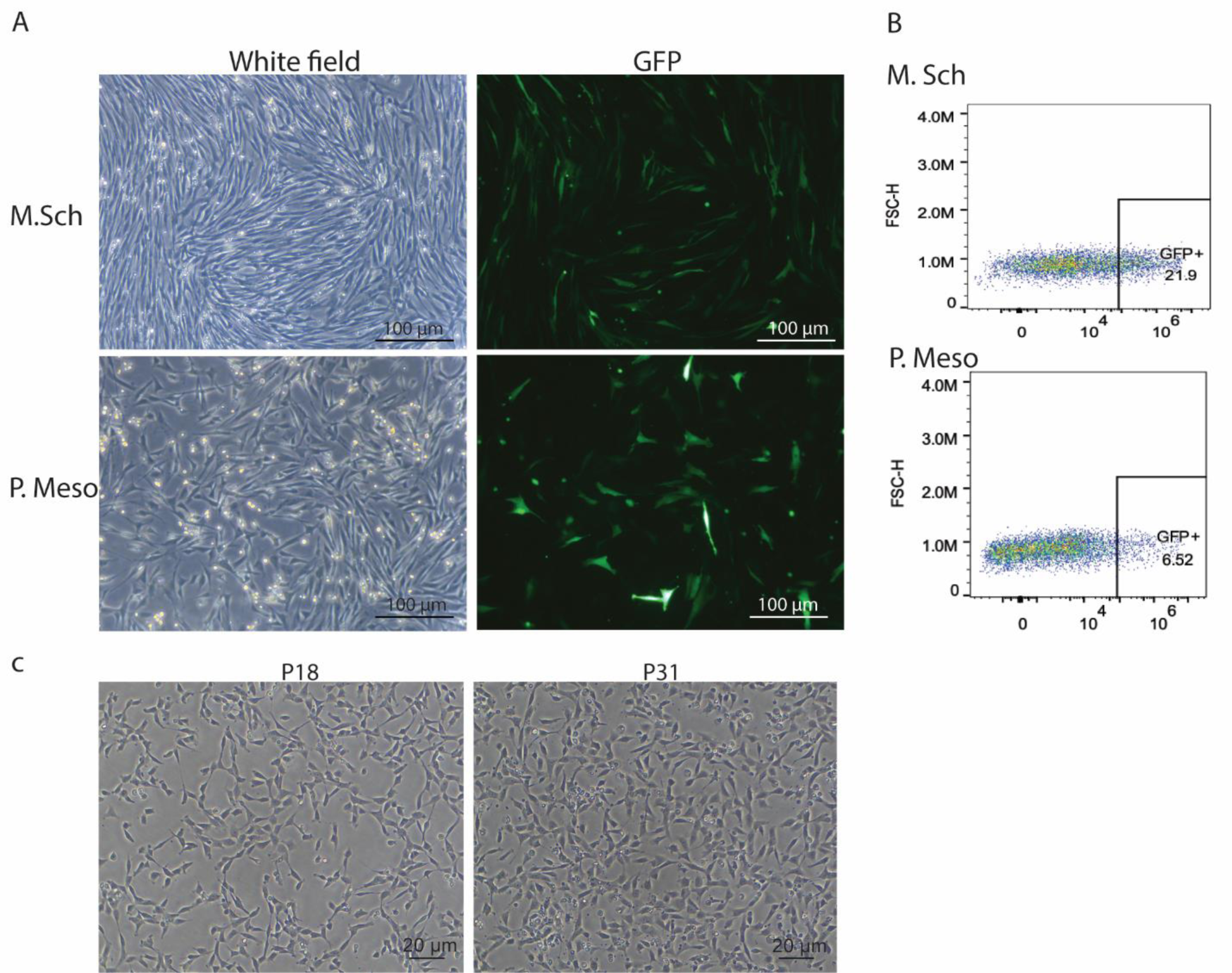
Lentiviral transduction of *M. Sch* and *P. Meso* fibroblasts. (**A**) white field and fluorescent images of lentiviral infected fibroblasts showing transduction efficiency of eGFP. (**B**) Representative flow cytometry plot of bat fibroblasts transduction efficiency. eGFP lentiviral infected bat fibroblasts were analyzed by Cytek® Aurora System.

**Figure 8.**
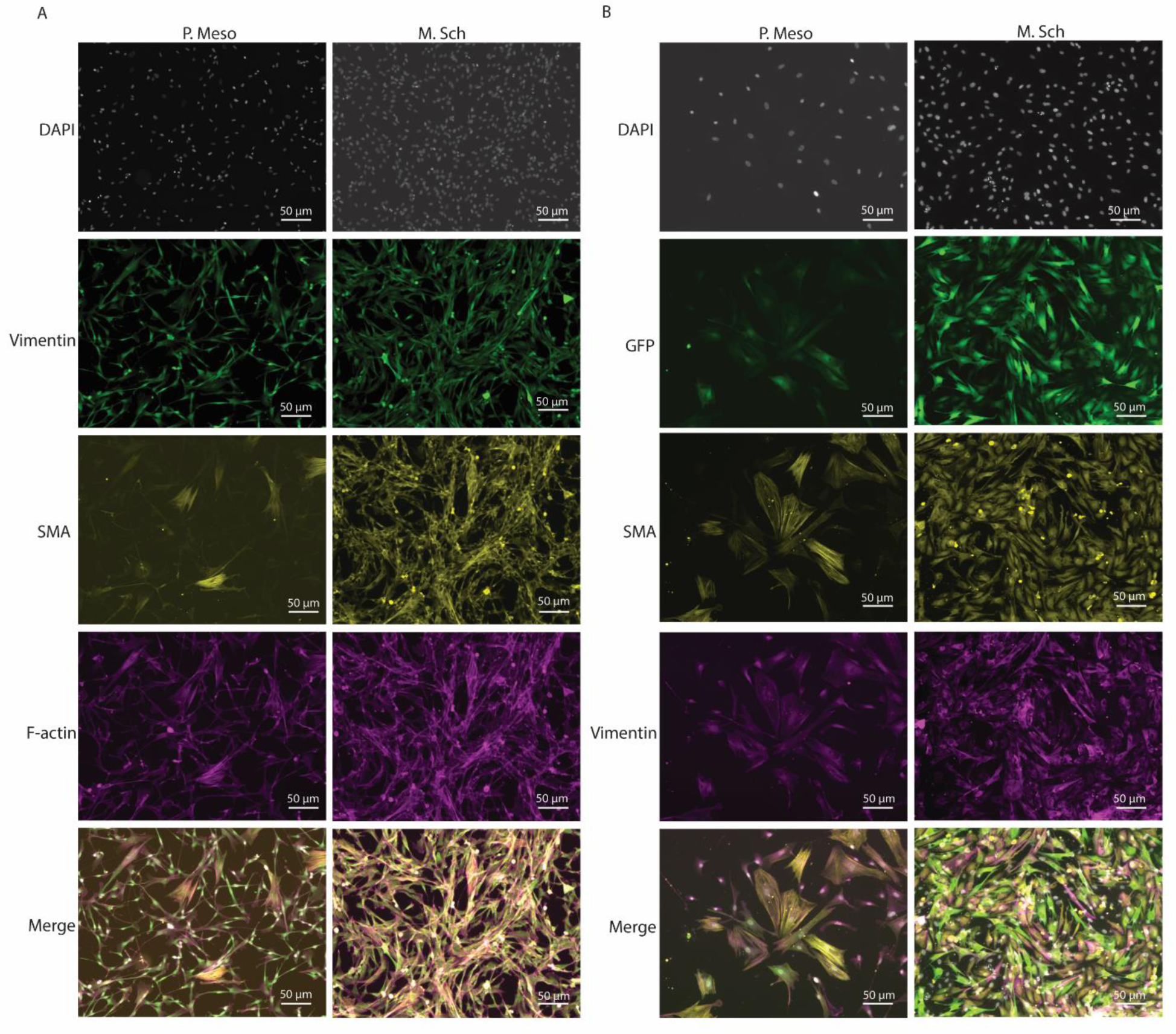
Immunofluorescence analysis of *M. Sch* and *P. Meso* primary fibroblasts and immortalized cell lines. A. Vimentin SMA and F-actin staining of *M. Sch* and *P. Meso* primary fibroblasts. Green: Vimentin; Yellow: SMA; Magenta: F-actin; Blue: DAPI. B. Vimentin and SMA staining of *M. Sch* fibroblasts *-TERT-GFP* and *P. Meso* fibroblasts -SV40-GFP/*hTERT-GFP.* Green: GFP; Yellow: SMA; Magenta: Vimentin; Blue: DAPI.

**Figure 9.**
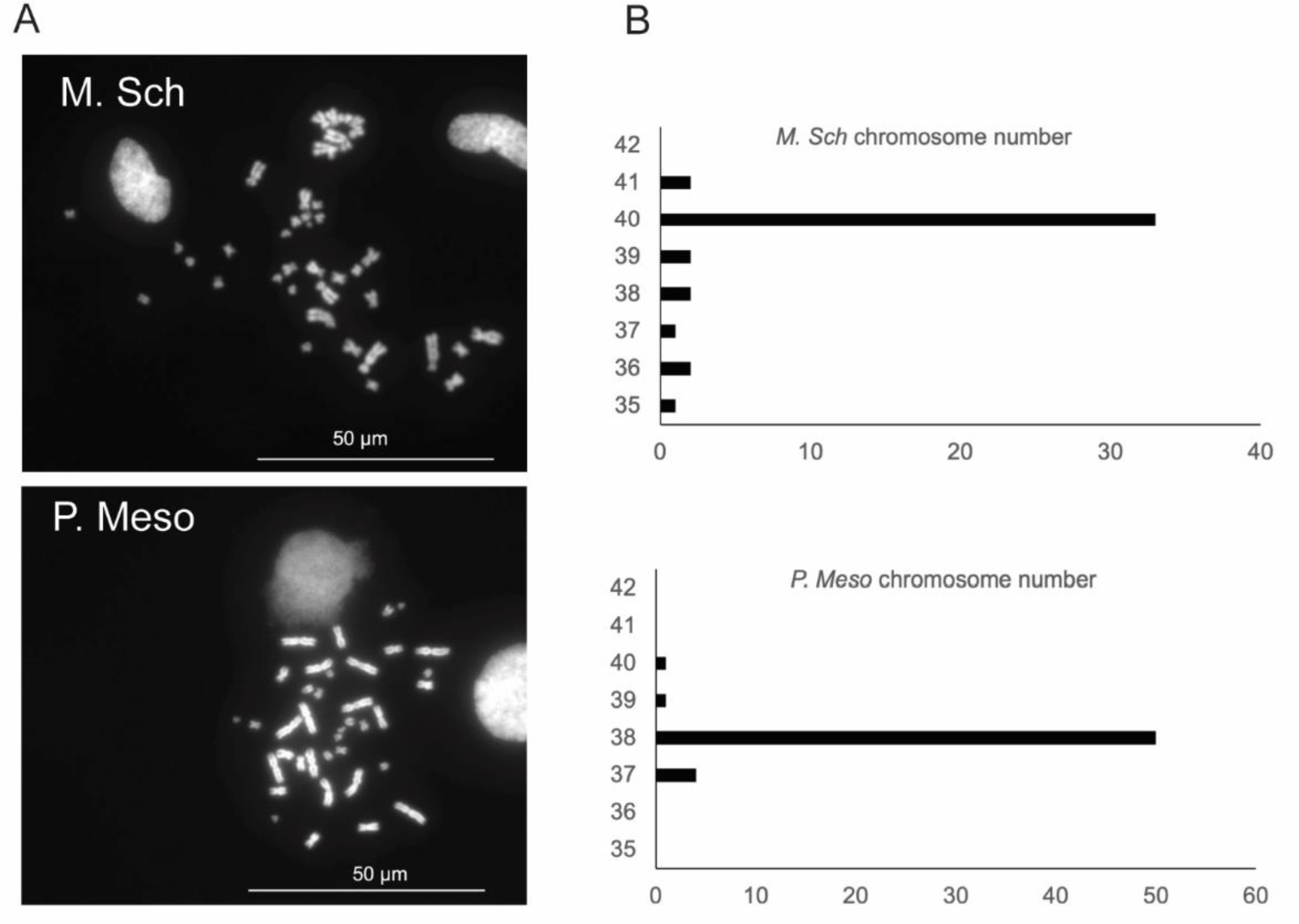
Karyotyping of *M. Sch* and *P. Meso* fibroblasts. A Representative chromosome spread for *M. Sch* and *P. Meso* fibroblasts. B. The graph shows the chromosome counting of *M. Sch* and *P. Meso* fibroblasts. The chromosome numbers of 43 and 55 metaphase chromosome spreads respectively for *M. Sch* and *P. Meso* fibroblasts were counted.

### Basic Protocol 1: Bat wing biopsy collection and preservation

*Note: The bat wing biopsy collection is conducted with a wildlife collecting license*.

*Note: Export and import permits must be obtained according to each country’s requirements*.

This protocol describes the detailed method to collect and preserve bat wing biopsies (Fig. 1)

## Materials Requirement

### Reagents, Solutions and Requirements

- Sterile 70% isopropyl alcohol prep pads (VWR, Cat#75840-802)
- Sterile povidone-iodine antiseptic swabsticks (Dynarex, Cat#1201)
- Ethanol, 70%
- Trypsin-EDTA (0.25%, STEMCELL Technologies, Cat# 07901)
- Dimethyl sulfoxide hydri-max (DMSO) (Sigma, Cat# 67-68-5)
- FBS (PEAK, Cat# PS-FB2)
- Liquid nitrogen (LN2)
- Cryovial (Corning, Cat# 430659)
- Mr. Frosty® Freezing Container (ThermoFisher Scientific, Cat# 5100-0001) or Corning^®^ CoolCell^™^ LX Cell Freezing Container (Sigma, Cat# CLS432004)
- Disposable surgical gloves
- Biopsy Punches 3 mm disposable (VWR, Cat# 21909-136)
- Sterile, single use petri dishes
- Forceps (Fine Science Tools, Cat# 91150-20)

1. Gently wrap bat with a cotton bag with their faces covered (Fig. 1A). *Note: Bats should be gently wrapped with a cotton bag with their faces covered to calm bats (7g-50g), to reduce stress and the risk of injury*.
2. Prior to biopsy, extend bat wing (Fig. 1B) with a back light to ensure major blood vessels are not present in biopsy area.
3. Sample the area(s) on the flight membrane closer to the body wall, which heal rapidly.
4. At sample site, cleanse the skin thoroughly using alcohol prep pads until the pad shows no traces of dirt (Fig. 1C).
5. Prepare the same site by applying antiseptic with a swab (Fig. 1C).
6. Extend and position the wing against a flat, hard surface with a sterile petri dish when using the biopsy tool. Keep biopsy punch perpendicular to the wing while pressing and twisting to cut out (Fig. 1C). *Note: Do not rock the tool side-to-side*.
7. Use sterile forceps to collect the 3 mm skin biopsy that remains behind on the cutting surface of the petri dish. If the biopsy is lodged in the biopsy punch, use forceps to remove the sample.
8. Rinse biopsy punch with antiseptic for 30 s and wash in 1x PBS to remove it (Fig. 1D).
9. Transfer clean tissue into a 2 mL centrifuge tube filled with 1 mL 0.05% EDTA-Trypsin and incubate for 40 min at ambient temperature (Fig. 1E).
10. To the same tube, add 1 mL Freezing Media 1 (Table 1) (Fig. 1E).
11. Immediately, place the cryovials into a Freezing Container on dry ice (-78.5° C) for a minimum of 80 minutes (-1° C per minute) before transferring to a -80° C freezer or LN2 (Fig. 1E).
12. Store in a -80° C freezer for up to 4 months or in LN2 for long-term storage until primary cell derivation.

### Support Protocol 1: Blood collection from bat-venipuncture

*Note: Bat derived cells should be obtained from healthy (rabies-free) bats. The health status can be screened from non-lethal blood serum collection for testing by the Kansas State Veterinary Diagnostic Laboratory (USA). However, if rabies-neutralizing antibodies are detected, the test cannot differentiate whether the immune response is from current infection or prior rabies virus exposure* [34].

**Table 1.** Freezing Media 1 (10 mL)

| Reagent | Volume | Final Concentration |
| --- | --- | --- |
| FBS | 8 mL | 80% |
| DMSO | 2 mL | 20% |

*Note: If lethal samples are obtained, the brainstem should be obtained, stored in* RNAlater™ Stabilization *Solution (ThermoFisher, AM7020) or flash frozen, and tested for rabies with RT-PCR* [35].

## Additional Materials

- VivaGuard® Lancet 30G (VWR, Cat# 76670-432)
- PTS Collect Capillary Tube 50μL (CLIAwaived, Cat# CHEK-2134)
- Screw-Cap Microcentrifuge Tubes 0.5mL (VWR, Cat# 10025-744) or MiniCollect® Serum Tubes (VWR, Cat# 95057-286)
- Low speed centrifuge
- Cryotubes 0.5ml (VWR, Cat# 76642-646)

1. For added protection of the handler, an anti-cut finger protector can be placed on the index finger of the non-dominant hand with a second glove to cover it.
2. The free wing of the bat (Fig. 1A, iv) should be outstretched, with the forearm resting between the thumb and index finger of the non-dominant hand (Fig. 1A, v). This extends the wing in an upstroke position enabling visualization and sampling from the cephalic or median antebrachial vein (Fig. 1B).
3. Using the dominant hand, a 30G lancet should be held at a 15–30-degree angle with the needle bevel facing up. Puncture the vein in a smooth quick motion.
4. Quickly collect blood using a 50 μL capillary tube and transfer to a 0.5 mL microcentrifuge tube. *Note: Limit blood collection to a maximum of 10% body weight. For a 10 g bat, collect a maximum of 100 μL blood sample*.
5. After blood collection, apply direct pressure to stop the bleeding.
6. The bat should be offered fluids and food to recover (Fig. 1A).
7. Place bat in a cotton bat bag and allow to rest for at least 5 min to ensure hemostasis has occurred.
8. Allow the blood sample to clot at ambient temperature 30 - 45 minutes.
9. Pellet blood clot by centrifugation at 1,000-1,500 xg for 10 minutes using a refrigerated centrifuge.
10. Following centrifugation, immediately transfer the serum into sterile 0.5 mL cryotubes. Flash freeze and save blood clot or store in RNALater for collaborative projects found at https://www.gbatnet.org/interdisciplinary-projects/
11. Store serum in LN2.

### Basic Protocol 2: Isolation of primary fibroblasts from adult bat frozen wing biopsy

This protocol describes the detailed method to isolate primary fibroblasts from bat frozen wing biopsy collection (Fig. 2).

*Note: The bat primary fibroblast isolation procedure and cell culture are all performed in a biosafety cabinet in a biological safety level 2 room*.

*Note: All solutions and dissection tools used in isolation of primary fibroblast processing as well as cell culture processing must be sterile*.

**Additional Materials** (other materials are included in basic protocol 1)

## Reagents, Solutions and Requirements

- Advanced DMEM/F-12 (Gibco, Cat# 12-634-010)
- GlutaMAX™ Supplement (Gibco, Cat# 35050061)
- HyClone Characterized FBS (Cytiva, Cat# SH30071.03)
- bFGF (Gibco, Cat# PHG0021)
- Penicillin/Streptomycin (P/S) (Gibco, Cat# 15140122)
- DPBS (Corning, Cat# 21040CV)
- Gelatin solution (Millipore, Cat# ES-006-B)
- Amphotericin B (R & D System, Cat# B23192)
- Dispase (1 U/mL) (STEMCELL Technology, Cat **#** 07923)
- Collagenase Type IV (STEMCELL Technology, Cat **#** 07427)
- DMEM (ATCC, Cat# 30-2002)
- FBS (PEAK, Cat# PS-FB2)

## Hardware and Instruments

- 6-well culture plate (Corning, Cat#3516)
- 12-well culture plate (Corning, Cat# 3513)
- Conical tubes (15 mL) (VWR, Cat# 21008-918)
- Portable pipette aid and pipette
- Micro pipettes (Integra)
- Inverted Microscope (Leica)
- Centrifuge (Thermo Scientific Heraeus Megafuge 8)
- 37°C Incubator
- Curved and straight scissors with sharp ends
- Disposable scalpel (Sklar Instruments, Cat# 06-3120)

1. Take out the frozen vial with wing biopsy from LN2, and transfer to BSL2 lab on dry ice.
2. Thaw frozen cryovials in hand and immediately transfer wing biopsy with tweezers into one well of a 6-well plate that is preloaded with 3 mL of DPBS+5X P/S and incubate for 5 min at RT. *Note: hereafter, all the cell derivation and culture experiments will be carried out in a biosafety cabinet in a BSL2 room*.
3. Transfer the wing biopsy to another new well of the same 6-well plate with 3 mL of DPBS+5X P/S, incubate for 5 min at RT.
4. Repeat step 3 one time.
5. Mince wing biopsy into small pieces with a sterile scalpel or scissor (Fig. 3A).
6. Add 3 mL of dispase solution to each well, ensure the tissue pieces are submerged in solution. Incubate at 37 °C for 90 min.
7. Pipet the tissue up and down about 5-10 times by 1000 µL pipet.
8. Observe under microscope. If the wing biopsy separated into 2 layers (Fig. 3B), go to step 9. If not, continue incubating in dispase solution at 37 °C for another 30 min.
9. Transfer the tissue into a new well of 6-well plate preloaded with 3 mL collagenase Type IV solution (1 mg/mL). Incubate at 37 °C for 12 h.
10. Pipet the tissue up and down about 10 times by 1000 µL pipet.
11. Observe under microscope. Most tissue should be dissociated into very small pieces or cell clusters (Fig. 3 C).
12. Add 3 mL bat primary fibroblasts culture media to stop collagenase IV dissociation.
13. Transfer cells into a 15 mL tube.
14. Centrifuge at 1000 rpm for 5 min.
15. Aspirate the supernatant, resuspend the cells in each tube with 1 mL bat primary fibroblasts culture media (Table 2).
16. Seed cells/small tissue pieces in 12-well culture plate (coated with 0.1% gelatin).
17. Incubate cells at 37 °C, with 5 % CO2. *Note: For culture of primary fibroblasts derived from adult bat frozen wing biopsy, 2-5 days after seeding, attached cells will be visible under microscope (Fig. 3D). Note: P0 cells might show different morphologies, indicating different cell types including fibroblasts and endothelial cells, epithelial cells (Fig. 3D)*.

## Support Protocol 2: Primary fibroblasts culture and subculture (Fig. 4)

1. After initial seeding of the primary culture, observe the cells under microscope every day until cells are 50-90% confluent (4-6 days).
2. Transfer the old media together with the small tissue pieces and suspension cells to another well of the 12-well plate. *Note: Primary fibroblasts will be continually released from the tissue explants and attach to the bottom of the tissue culture dish (Fig. 4C, D, E, F)*. *Note: We kept transfer old media together with the small tissue pieces and suspension cells to new wells at least 3 times for primary cell derivation.* *Note: With this method, the bat wing tissue explants were able to survive for >100 days and keep releasing cells in the bat primary fibroblasts culture media (Fig. 4G, H)*.
3. Wash cells once with bat primary fibroblasts culture media and transfer the supernatant to the same well in step 2.
4. Add 300 µL of TrypLE to each well (12-well plate), incubate 5 min at 37 °C.
5. Add 300 µL bat primary fibroblasts culture media to stop the dissociation.
6. Transfer cells to a 15 mL tube.
7. Centrifuge at 1000 rpm for 5 min.
8. Aspirate the supernatant, resuspend the cells with 1 mL bat primary fibroblasts culture media.
9. To expand cells, plate the cells from 1 well of 12-well plate into 2 wells of a 6-well plate coated with 0.1% gelatin. Incubate cells at 37 °C, with 5 % CO2. *Note: Hereafter, we can keep expanding the cells.* *Note: For the 2 primary bat wing fibroblasts from 2 different bat species P. Meso and M. Sch, we usually subculture them at 1:4 ratio, every 3-4 days.* *Note: fibroblasts derived from bat frozen wing biopsy can be cultured in bat primary fibroblasts culture media for more than 20 passages without observable morphological changes (Fig. 4A, B).* *Note: after 3 passages, most cells showed spindle shaped or stellate shaped fibroblasts morphology*.

## Support Protocol 3: Growth curve and doubling time

In cell culture, growth curves and doubling time are critical parameters that provide valuable insights into the behavior and characteristics of cultured cells.

1. Seed bat fibroblasts (different passages) in 1 mL cell culture media (normal fibroblasts media (Table 3) or bat primary fibroblasts culture media (Table 2)) in wells of 12-well plates at a concentration of 4-8 e^4^ cells/well. Incubate cells at 37 °C, with 5 % CO2. *Note: Culture bat primary fibroblasts in bat primary fibroblasts culture media is critical for bat primary fibroblasts derivation (Fig. 5).* *Note: Bat primary fibroblasts represent different morphologies when cultured in different media (Fig. 5A, B)*.
2. Take images (Fig. 6A) and count cell numbers every day (Fig. 6B). Calculate cell doubling time through: https://www.doubling-time.com/compute.php (Fig. 6C) *Note: the cells growth curve and doubling time for both bat species are consistent among early passages <P11 when using bat primary fibroblasts culture media (Fig. 6).* *Note: the cells growth curve and doubling time for M. Sch fibroblasts cultured in different media are different (Fig. 5C)*.

**Table 2.** Bat primary fibroblasts culture media composition (100 mL)

| Reagent | Volume | Final Concentration |
| --- | --- | --- |
| Advanced DMEM/F-12 | 87.59 mL | 90% |
| HyClone Characterized FBS | 10 mL | 10% |
| Glutmax | 1 mL | 1% |
| bFGF (0.1 mg/mL) | 0.01mL | 10 ng/mL |
| Amphotericin b | 0.4 mL | 1 µg/mL |
| Penicillin/Streptomycin | 1 mL | 1X |

**Table 3.**
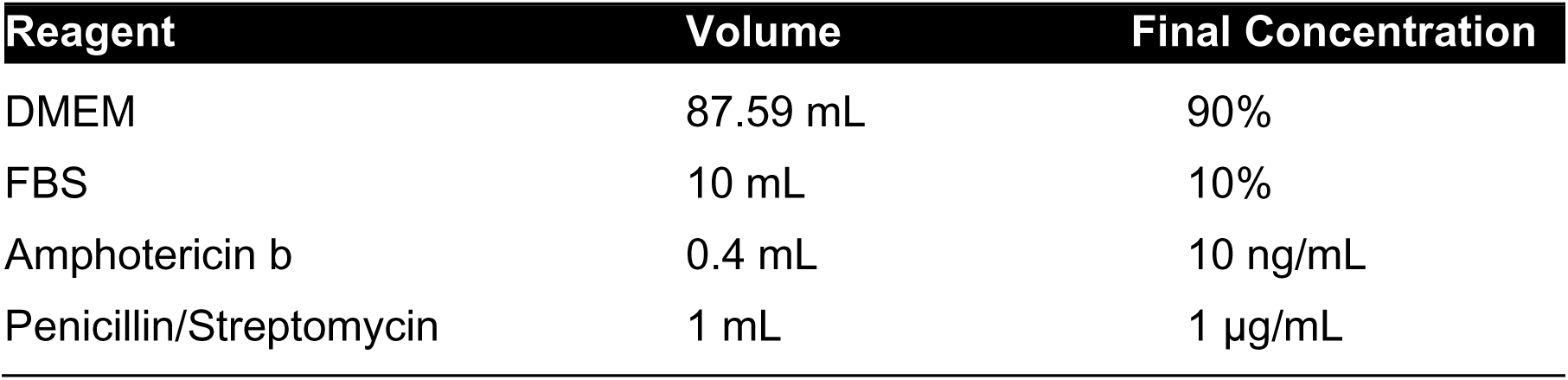
Normal fibroblasts media composition (100 mL)

| Reagent | Volume | Final Concentration |
| --- | --- | --- |
| DMEM | 87.59 mL | 90% |
| FBS | 10 mL | 10% |
| Amphotericin b | 0.4 mL | 10 ng/mL |
| Penicillin/Streptomycin | 1 mL | 1 µg/mL |

## Support Protocol 4: Cell banking and thawing of primary fibroblasts

*Note: We recommend cryopreserving bat primary fibroblasts at P3, P4 or P5 when the culture is approximately 90% confluent and actively growing*.

*Note: Make fresh freezing media each time before the cryopreservation of cells*.

## Cell banking

1. Aspirate the media from the culture plate (6-well plate) and wash the cells once with 2 mL DPBS.
2. Add 1 mL TrypLE, and incubate for 5 min at 37 °C.
3. Add 1 mL bat primary fibroblasts culture media.
4. Transfer cells to a 15 mL tube.
5. Centrifuge the cells at 1000 rpm for 5 min.
6. Remove supernatant and resuspend the cells in freezing media 2 (Table 4) (1 mL/well of 6-well plate)
7. Transfer cells into cryovials at 1 mL/vial (1 vial/well of 6-well plate).
8. Immediately, place the cryovials into a Mr. Frosty® Freezing Container, place at -70 °C, overnight.

*Note: Cryovials must be moved to a liquid nitrogen vapor tank for long-term storage*.

**Table 4.** Freezing Media 2 (10 mL)

| Reagent | Volume | Final Concentration |
| --- | --- | --- |
| HyClone FBS | 9 mL | 90% |
| DMSO | 1 mL | 10% |

*Note: Make fresh freezing media each time before the cryopreservation of cells.*

## Thawing of primary fibroblasts

9. Thaw 1 vial of frozen cells in hand or in 37 °C bead bath.
10. Immediately transfer the cells to a 15 mL tube with 9 mL pre-warmed bat primary fibroblasts culture media.
11. Centrifuge the cells at 1000 rpm for 5 min at RT.
12. Remove the supernatant, resuspend the cell pellet in 10 mL of pre-warmed bat primary fibroblasts culture media.
13. Seed the cells in a T25 cell culture flask. Incubate cells at 37 °C, with 5 % CO2.

## Support Protocol 5: Lentiviral transduction of bat primary fibroblasts

Genome editing is an important tool for gene functional analysis and genetic therapies. However, genome editing in low-passage primary cells is historically difficult due to their high physiological relevance to in vivo cells. Lentiviral particles are well known for their ability to deliver genetic molecules efficiently and stably into a variety of target cells. To test the possibility of whether bat primary fibroblasts can be genetically modified, we produced lentiviral particles and transduced the bat wing primary fibroblasts. Fig. 7 represents the lentiviral transduction of bat wing primary fibroblasts.

**Additional Materials** (other materials are included in basic protocol 1 and 2)

Lipofectamine™ 3000 Transfection Reagent (Invitrogen TM, Cat# L3000001)

Opti-MEM Reduced serum media (ThermoFisher Scientific, Cat# 31985062)

pIB2-SEC13-mEGFP (gifted from Dr. Subu Ramanathan)

VSV-G (Cat # 12259-pMD2.G)

psPAX2 (Cat # 12260-psPAX2)

Polybrene (R&D Systems, Cat# 7711)

0.45 µM filter (Sigma-Aldrich, Cat# SE1M003M00)

Dulbecco’s modified Eagle’s media (DMEM) (ATCC, Cat # 30-2002)

Sodium Pyruvate (100mM) (Corning, Cat# 25-000-CI)

## Lentivirus production

### Day 1: Seed 293T cells

1. Plate 293T cells at 2.0e^6^ cells per well in a 6-well plate in DMEM containing 10% FBS, 1x GlutaMax and incubated overnight at 37°C with 5% CO2 (90-99% confluency at time of transfection).

### Day 2: Transfection

2. Bring out Opti-MEM reduced serum medium to room temperature.
3. On the early morning of lentivirus packaging day, aspirate the media from the 293T cells culture dish and add 2 mL lentivirus packaging media (pre-warmed) (Table 5).
4. Add 7 μL Lipofectamine 3000 Transfection Reagent to a microcentrifuge tube with 250 μL Opti-MEM Reduced serum medium in a 1.5 mL tube. Mix well.
5. Add 1.25 µg of lentiviral expression plasmid pIB2-SEC13-mEGFP, 1 µg of psPAX2 and 0.25 µg pMD2.G, 6 μL P3000 Enhancer reagent to another 1.5 mL microcentrifuge tube with 250 μL Opti-MEM reduced serum medium. Mix well.
6. Prepare lipid-DNA complexes by adding the solution from step 4 to the solution from step 5. Mix well. Incubate at RT for 15 min.
7. Remove 1 mL of the lentivirus packaging media from each 293T culture well, leaving 1 mL in each well.
8. Add the 500 µL lipid-DNA complexes dropwise to the cells.
9. Incubate the cells at 37°C with 5% CO2 for 6 hours.
10. Carefully aspirate the media containing lipid-DNA complexes from the 293T cells culture dish and add 3 mL bat primary fibroblasts culture media without Amphotericin b, or Penicillin/Streptomycin (P/S) (pre-warmed).
11. Incubate the cells at 37°C with 5% CO2.

*Note: bat primary fibroblasts culture media without Amphotericin b, or P/S was used to collect lentivirus to avoid the stress from media during the transduction step*.

**Table 5.** Lentivirus packaging media composition.

| Reagent | Volume | Final Concentration |
| --- | --- | --- |
| Opti-MEM Reduced serum medium | 47.4 mL | 95% |
| HyClone FBS | 2.5 mL | 5% |
| Sodium Pyruvate (100mM) | 0.1 mL | 20 μM |

### Day 4: first batch of lentivirus collection

1. At 48 hours post transfection, collect the 3 mL media containing virus from each well into a 15 mL tube and store at 4°C.
2. Add 3 mL of fresh bat primary fibroblasts culture media without Amphotericin b, or Penicillin/Streptomycin (pre-warmed) to each well.
3. Incubate the cells at 37°C with 5% CO2.

### Day 5: second batch of lentivirus collection

4. At 72 hours post transfection, collect the 3 mL viral supernatant from each well and combine with the first collection.
5. Filter through a 0.45 µM filter, and store at 4°C for up to 1 week.

*Note: For long term storage, aliquot virus, and store at -80°C*.

## Lentiviral transduction of bat primary fibroblasts

### Day 0. Seed cells

1. Seed cells in a 12-well plate in 1 mL bat primary fibroblasts culture media at an appropriate density (60-75% confluency at time of transduction).
2. Incubate the cells at 37°C with 5% CO2 overnight.

### Day 1. EGFP Lentiviral transduction of bat primary fibroblasts

3. Aspirate the media from the culture dish (12-well).
4. Add 1 mL GFP lentivirus supernatant and 2 μl 4 mg/mL polybrene to the cells (polybrene working concentration 8 ug/ml).
5. Centrifuge the cells at 1000 g for 2 hrs at RT.
6. Transfer cells to cell culture incubator (37°C with 5% CO2).
7. At 24 hours post transduction, replace the media with fresh bat primary fibroblasts culture media.
8. Check the transduction efficiency under florescent microscope at 2 days post transduction (Fig. 7A) and sort cells by cytometry to count GFP positive cells (Fig. 7B)

## Basic Protocol 3: Bat stable fibroblasts cell lines development

One of the major issues of primary cell culture is their limited proliferation ability in culture [36], which can be a challenge for long-term experiments that require large number of cells. Immortalizing cell lines is a common strategy to overcome the limited lifespan of primary cells. There are several strategies that can be applied to immortalize cell lines. One of the important strategies is stable expression of Simian virus 40 T-antigen (SV40T), which would promote DNA replication and host cell division through down regulating the tumor suppressor proteins [37, 38]. Another commonly employed strategy is stable expression of the catalytic subunit of the human telomerase reverse transcriptase (hTERT), which is an enzyme that maintains telomere length, preventing cell senescence [39, 40]. Here, we describe bat fibroblasts immortalization through introduction and stable expression of hTERT, SV40 large T-antigen or both under CMV promoter.

**Additional Materials** (other materials are included in basic protocol 1 and 2)

hTERT (CMV, GFP-Puro) lentivirus in PBS (GenTarget Inc, Cat#: LVP1130-GP-PBS)

SV40 large T-antigen (CMV, GFP-Puro) lentivirus in PBS (GenTarget Inc, Cat#: LVP016-GP)

### Day 0. Seed cells

1. Seed cells in a 24-well plate in 0.5 mL bat primary fibroblasts culture media at an appropriate density (60-75% confluency at time of transduction). *Note: passage 3 cells of both species were used for immortalization*.
2. Incubate the cells at 37°C with 5% CO2.

### Day 1. hTERT and SV40 Large T-antigen Lentiviral transduction of bat primary fibroblasts

3. Thaw hTERT and SV40 large T-antigen Lentiviral at RT.
4. Add 50 µL hTERT lentiviral (or 50 ul SV40 large T-antigen Lentiviral, or 50 μL hTERT and SV40 large T-antigen lentiviral) to each well of the 24-well plate cultured with bat primary fibroblasts.
5. Centrifuge the cells at 1000g for 2 hrs at RT.
6. Transfer cells to cell culture incubator (37°C with 5% CO2).
7. At 24 hours post transduction, replace the media with fresh bat primary fibroblasts culture media. Incubate the cells at 37°C with 5% CO2.

### Day 3. Start selection by adding puromycin

8. At 48 hours post transduction, replace the media with fresh bat primary fibroblasts culture media with 2 ug/mL puromycin. *Note: kill curved assay should be done to decide the puromycin concentration before selection*.
9. Change media with 2 ug/mL puromycin every other day.
10. Subculture Cells when the culture is approximately 90% confluent.
11. Confirm fibroblasts cell line after 2 weeks of puromycin selection by IF staining and fluorescent microscope imaging. *Note: One bat cell line (P. Meso-hTERT/SV40-GFP) developed through hTERT/*SV40 *Large T-antigen transduction has been cultured for more than 30 passages*.

## Support Protocol 6: Bat fibroblasts validation by immunofluorescence staining (IF staining)

The bat primary fibroblasts derivate from the frozen wing biopsy and the immortalized fibroblasts line can be characterized through IF staining to confirm fibroblasts specific markers, such as alpha smooth muscle actin (alpha-SMA) and vimentin. Here, we describe IF staining for bat fibroblasts validation.

**Additional Materials** (other materials are included in basic protocol 1 and 2)

4% paraformaldehyde (PFA) solution in PBS

PBS

Triton X-100 (Thermo Fisher Scientific, cat. no. 85111)

Anti-Vimentin (Millipore Sigma, Cat# AB1620) (used at 1:40)

Alexa Fluor^TM^ 647 Phalloidin (ThermoFisher, Cat# A22287) (used at 1:40) Anti-SMA (Agilent, Cat# 76542) (used at 1:50)

Donkey anti-goat IgG (Biotium, Cat# 20016) (used at 1:500) Donkey anti-mouse IgG (Biotium, Cat# 20105) (used at 1:500) DAPI (Biolegend, Cat # 422801) (used at 1:500)

ProLong™ Gold Antifade Mountant (ThermoFisher, Cat# P10144) Round cover slides

1. Seed cells on cover slides in 12-well plate.
2. Incubate the cells at 37°C with 5% CO2, until cells reach 70-90% confluency.
3. Remove supernatant, and rinse cells with PBS one time.
4. Add 4% PFA to cells, and incubate at RT for 30 min.
5. Remove 4% PFA and add 1 mL PBS buffer with 0.1% Triton X-100 (PBST), incubate 5 min at RT.
6. Remove PBST, and add fresh PBST, incubate 45 min at RT for permeabilization.
7. Remove PBST and add 1 mL superblock buffer. Incubate 45 min at RT.
8. Add Primary antibodies. Incubate at 4 °C overnight.
9. Rinse the cells with PBST, 5×5 min.
10. Apply secondary antibodies and DAPI to the cells and incubate in dark at RT for 1 hour.
11. Rinse well with PBST for 5×5 min.
12. Replace PBST with PBS.
13. Mount the cells with ProLong™ Gold Antifade Mountant.
14. Proceed for imaging. (Fig. 8) *Note: IF staining results showed that all the p3 primary cells and cell lines developed in this protocol express fibroblasts specific markers (Fig. 8)*.

## Support Protocol 7: Chromosome counting

Chromosome counting, also known as karyotyping, is a crucial technique in genetics and cytogenetics. In cell culture, it is a valuable tool for ensuring the authenticity, stability, and genetic integrity of cell lines. Here we describe the chromosome counting method to verify the bat primary fibroblasts genetic integrity.

**Additional Materials** (other materials are included in basic protocol 1)

Colcemid (10 ug/mL stock; Invitrogen Life Technologies, Cat# 15212-012).

Methanol (VWR, Cat# PL230ZA-4P)

Glacial acetic acid (Fisher Scientific, Cat# AC22214-0010)

KaryoMAX™ Potassium Chloride Solution (ThermoFisher, Cat # 10575090)

VECTASHIELD® Antifade Mounting Medium with DAPI (Vector, Cat# H-1200-10)

**Table 6.** Fixative (80 mL)

| Reagent | Volume | Final Concentration |
| --- | --- | --- |
| Methanol | 60 mL | 66.666% |
| glacial acetic acid Sodium | 20 mL | 33.333% |

*Note: Need to prepare fresh Fixative each time before experiment.*

*Note: Need to prepare fresh Fixative each time before experiment*.

1. Culture bat primary fibroblasts in bat primary fibroblasts culture media in T25 cell culture flask at 37°C with 5% CO2, until cells reach 50-60% confluency.
2. Add 50 μL colcemid (working concentration 50 ng/ml) to the cells for 3 hrs. Note: Add colcemid to the same growth media, do not change the media.
3. Remove the media.
4. Wash with PBS.
5. Add 3 mL TrypLE, and incubate for 5 min at 37 °C.
6. Add 3 mL bat primary fibroblasts culture media to stop dissociation.
7. Transfer cells to a 15 mL tube.
8. Centrifuge at 1000 rpm for 5 min.
9. Aspirate the supernatant, resuspend the cells with 1.5 mL bat primary fibroblasts culture media.
10. Add 10 mL of warmed (37 °C) KCL solution slowly, drop by drop, along the wall of the tube flicking the tube gently at the same time. Invert serval times.
11. Incubate at 37 °C for 40 min.
12. Add two or three drops of ice-cold fixative, invert the tube to mix.
13. Centrifuge at 1000 rpm for 5 min.
14. Aspirate the supernatant, leaving 1mL behind.
15. Gently suspend cells.
16. Add up to 5-10 mL of ice-cold fixative drop by drop and disperse the cells thoroughly in the fixative.
17. Centrifuge at 1000 rpm for 5 min.
18. Aspirate the supernatant, leaving 1mL behind.
19. Gently suspend cells.
20. Repeat step 16-19 at least three times.
21. Resuspend the cells in 1-2 mL fixative.
22. Drop the cell suspension (20 μL) onto a glass slide from a 10-cm distance.
23. Air-dry the slides,
24. Mount the slides with VECTASHIELD® Antifade Mounting Medium with DAPI and seal the slide with a coverslip.
25. Proceed for imaging under florescent microscope (Fig. 9 A).
26. Count the number of chromosomes in each spread (Fig. 9 B).

*Note: At least 40 good spreads should be counted and analyzed*.

## Commentary

### Background Information

Fibroblasts are a type of connective tissue cells that maintain the structural integrity of all organs [26]. They are responsible for producing and maintaining the ECM, which provides support and structure to surrounding cells. Fibroblasts play crucial roles in wound healing, inflammation, tissue regeneration, angiogenesis, cancer progression, pathological tissue fibrosis, and the advancement of virus infections [26]. Lineage tracing and multiomics single-cell analyses have revealed that fibroblasts are diverse and retain an embryonic gene expression signature inherited from their organ of origin, offering new prospects for tissue-specific antifibrotic therapies [41]. Given the significant roles of fibroblasts in various physiological and pathological processes, as well as their ease to work with, they have been utilized in numerous fields of research [41–44]. For example, Mitra et al. established an in vitro model of cellular quiescence in primary human dermal fibroblasts to study various disease states [45]. Alsharabasy et al. developeded a protocol to create a skin fibrosis model using human dermal fibroblasts. This model can be employed for investigating the possible anti-fibrotic properties of specific chemical compounds [46]. Recently, human in vitro skin models were also dveloped for wound healing and wound healing disorders [47]. Additionally, fibroblasts have also been used to study the mechanisms of host adaptation to viral infection [42].

Isolating high-quality and a large number of primary fibroblasts is essential for developing in vitro fibroblast models. Various tissue dissociation methods are employed for fibroblast isolation from different tissues [41, 45, 48–57]. For isolation of primary fibroblasts from the skin dermis, multiple protocols have been developed with each differing in enzymatic digestions and mechanical dissociation [50, 52, 55]. Yohe et al. described a protocol for primary fibroblast isolation from fresh bat wing biopsies. They used collagenase IV to dissociate the skin and were able to observe attached cells approximately 24 hours after cell plating [58]. However, this protocol did not work for isolating primary fibroblasts from frozen bat wing biopsy (this study), where cell integrity and viability are lower than in fresh tissue. Therefore, developing or optimizing a robust, standardized, high-yield protocol for isolation of primary fibroblasts from frozen bat wing biopsy is necessary for bat and other wild animal studies.

### Critical Parameters

By following this protocol, we have been able to successfully isolate, culture and expand primary fibroblasts from adult bat frozen wing biopsy of two bat species: *P.meso* and *M. sch*. The size of the frozen wing biopsy, freezing method, bat age, sex, and species might affect the time of primary cell release from tissue explants. For primary fibroblast isolation from *P. meso* frozen wing biopsy, attached cells were observed at 2 days post cell derivation, while it took 5 days to detect the attached cells for *M. sch.* The frozen tissue size from *P. meso* wing biopsy was about 3×3 mm before cryopreservation (non-lethal), while *M. sch* wing biopsy was cryopreserved as 1×1 cm (lethal sample). Despite the difference in the time required to obtain the primary cells, we were able to obtain a large number of cells from both species, indicating that the tissue itself might be an important variable in this difference.

The bat wing web skin is highly folded and comprised of two epidermal layers separated by a central core of dermal connective tissue [59]. Fibroblasts are embedded in the dermal connective tissue (Fig. 1A). Thus, splitting the epidermis from the dermis is critical to exposure dermal connective tissue containing fibroblasts for further step fibroblasts isolation. Dispase is a type of matrix metalloprotease which can split epidermal and dermal sheets [52, 60, 61]. And we found that it is a critical step for successful isolation of primary fibroblasts from bat frozen wing biopsy. Without dispase digestion, little cells were detected after 2 weeks, while it only takes 2-5 days to observe primary cells sprouting from the biopsy tissue.

Cell culture media plays a crucial role in primary cell culture, influencing cell growth, viability, and functionality [54]. Advanced DMEM/F-12 was employed for bat fibroblasts culture in this protocol, as it contains ethanolamine, glutathione, ascorbic acid, insulin, transferrin, AlbuMAX™ II lipid-rich bovine serum albumin, and trace elements including sodium selenite, ammonium metavanadate, cupric sulfate, manganous chloride, as well as non-essential amino acids. These compounds are thought to benefit fibroblasts in culture and maintain their proliferation and functionality. Basic fibroblast growth factor (bFGF or FGF-2) was also added to bat fibroblasts culture media to promote fibroblasts viability and proliferation. Comparing the growth curve and doubling time of bat primary fibroblasts cultured in bat fibroblast culture media and normal fibroblast media indicates that bat fibroblast culture media had a significant impact on the proliferation ability of bat primary fibroblasts. (Fig. 5). So, bat fibroblast culture media might be another key factor for the successful derivation of primary fibroblasts from frozen wing biopsies. With the bat fibroblast culture media, the pieces of wing tissue explants were able to survive for more than 100 days and continue releasing cells (Fig. 4), indicating that bat fibroblast culture media can maintain cell viability for long time.

### Troubleshooting Guide for derivation of bat primary fibroblasts from adult bat frozen wing biopsy and stable cell lines development

Common problems, possible causes, and suggested solutions are listed in Table 7.

**Table 7.** Troubleshooting Commonly Encountered Problems.

| Problem | Possible cause | Solution |
| --- | --- | --- |
| Cells not attached | Low cell viability<br>Bad tissue dissociation<br>Non-treated vessel surface | Add Rock inhibitor Y-27632 (10 μM) to increase the viability.<br>Coat the culture vessel surface with 0.1% gelatin.<br>Use dispase to split epidermal and dermal sheets. |
| Low cell viability and proliferation | Tissue freezing problem<br>Unsuitable cell culture media | Cut wing biopsy into small pieces before freezing the tissue.<br>Use good dissociation method.<br>Use bat fibroblasts culture media. |
| Some samples grow slower than others. | sample-to-sample differences<br>sample size | Subculture cells when the cells reach >70% confluency.<br>Use enough tissue for primary cell derivation. |
| Low transduction efficiency | Low lentivirus packaging efficiency<br>Transduction method | Packaging lentivirus following the protocol.<br>Decrease cell number while increase lentivirus amount.<br>Use spinoculation transduction protocol |

### Understanding Results

By following basic protocol 2, healthy and actively proliferating bat primary fibroblasts can be obtained within 2 weeks. The time to obtain enough number of primary fibroblasts might be different due to tissue size, bat species, or ages. This protocol isolates the different cell populations from the bat wing biopsy (Fig. 3D). Different type of cells can be detected at P0 (Fig. 3D). With bat primary fibroblasts media, fibroblasts become enriched during the cell culture. The results of fibroblast molecular marker staining indicate that all primary cells at P3 are fibroblasts (Fig. 8A).

Bat primary fibroblasts grow fast. More than 20e6 cells can be obtained within 3 weeks from 1 cm X 1 cm frozen wing biopsy. We found that bat wing explants cultured in the bat fibroblast culture media were able to survive for more than 100 days and continually release cells. This might be due to the suitability of the cell culture conditions for the bat wing explant or the special characteristics of the bat wing tissue. Both *P. Meso* and *M. Sch* primary fibroblasts cell lines were able to be propagated healthily for more than 20 passages. This is surprising because primary fibroblasts from other mammals, such as mouse or human, can only be subcultured for a limited time (usually < 10 passages). These results might indicate that our fibroblast cell culture conditions are conducive to fibroblast viability and proliferation, or that bat fibroblasts differ from those of other mammals and auto-immortalize. Up to date, the immortalized cells have been propagated for # passages.

### Time Considerations

The procedure timeline estimation is detailed below:

Bat wing biopsy collection and cryopreservation (approx. 2 hours per bat)

Bat frozen wing biopsy dispase digestion (approx. 90 min)

Collagenase IV dissociation (approx. 12-14 hours) Trypsin-EDTA dissociation (approx. 15 min)

Cell collection (approx. 10 min)

Primary fibroblast cell cultures (approx. 5 to 10 days)

Bat stable fibroblasts cell lines development (>3 weeks)

IF staining (approx. 2 days)

## Acknowledgments

The fieldwork was carried out within the framework of the informally dubbed “Belize Bat-a-thon”, founded by Dr. Brock Fenton and co-organized by Dr. Nancy Simmons and we thank everyone for the field support. The authors would like to thank the Stowers Institute for Medical Research technology centers: Cells, Tissues, and Organoids Center members for the meaningful discussion throughout protocol optimization; Media Preparation Center for preparing media. We also thank Scientific Illustrator Mark Miller for creating the bat wing structure image and the overview of bat wing fibroblasts derivation in Figure 2. We acknowledge that the pIB2-SEC13-mEGFP plasmid was gifted from Dr. Subu Ramanathan. This work was supported by NIH grant 1DP2AG071466-01 to NR for the field study. Supplemental funding was provided by the NSF Postdoctoral Research Fellowships in Biology #2109717 to J.C., the BWF PDEP grant (G-1022339) to J.C., and the HHMI Hanna H. Gray Fellows Program (GT15991) to J.C. Any opinions, findings, and conclusions or recommendations expressed in this material are those of the authors and do not necessarily reflect the views of the NSF, BWF, or HHMI.

## Author contributions

Fengyan Deng: study design, methodology, data curation, data validation, writing, figure design; Pedro Morales-Sosa: data validation; Andrea Bernal-Rivera: field study, Yan Wang: manuscript editing; Dai Tsuchiya: data validation; Jose Emmanuel Javier: data validation, Nicolas Rohner: manuscript editing; Jasmin Camacho: conceptualization, study design, field study, manuscript editing, figure design; Chongbei Zhao: conceptualization, investigation, supervision, writing.

## Data Availability Statement

Data sharing not applicable to this article as no datasets were generated or analyzed during the current study.

